# Energy partitioning in the cell cortex

**DOI:** 10.1101/2024.05.06.592707

**Authors:** Sheng Chen, Daniel S. Seara, Ani Michaud, Songeun Kim, William M. Bement, Michael P. Murrell

**Affiliations:** Department of Biomedical Engineering, Yale University, New Haven, Connecticut 06511, USA; Department of Physics, Yale University, New Haven, Connecticut 06511, USA; Systems Biology Institute, Yale University, West Haven, Connecticut 06516, USA; James Franck Institute, University of Chicago, Chicago, IL, USA; Cellular and Molecular Biology Graduate Program, University of Wisconsin-Madison, Madison, WI, USA; Center for Quantitative Cell Imaging, University of Wisconsin-Madison, Madison, WI, USA; Department of Integrative Biology, University of Wisconsin-Madison, Madison, WI, USA

**Author notes:** Correspondence should be addressed to M.P.M. or W.M.B.,). These two authors contributed equally to this work (William M. Bement, Michael P. Murrell).

## Abstract

Living systems are driven far from thermodynamic equilibrium through the continuous consumption of ambient energy^1^. In the cell cortex, this energy is invested in the formation of diverse patterns in chemical and mechanical activities, whose unique spatial and temporal dynamics determine cell phenotypes and behaviors^2-6^. However, how cells partition internal energy between chemical and mechanical work is unknown^7-9^. Here we measured the entropy production rate (EPR) of both the chemical and mechanical subsystems of the cell cortex across a broad range of periodic patterns as the system is driven further from equilibrium via manipulation of the Rho GTPase pathway, which controls cortical actin filaments (F-actin) and myosin-II. We find that at lower levels of Rho GAP (GTPase activating protein) expression, which produce pulses or “choppy” Rho and F-actin waves, energy is comparably partitioned between the chemical and mechanical subsystems and is subject to the constraint of Onsager reciprocity. Within the range of reciprocity, the EPR is maximized in choppy waves that resemble the waves associated with cell division^3,10^. However, as the cortex is driven even further from equilibrium into elaborate labyrinthine or spiral traveling wave trains via increased GAP expression, reciprocity is broken, marking an increasingly differential partitioning of energy and an uncoupling of chemical and mechanical activities. We further demonstrate that energy partitioning and reciprocity are determined by the competition between the timescales of chemical reaction and mechanical relaxation. These results indicate that even within coupled cellular subsystems, both the relative proportions of energy partitioned to each subsystem and the ultimate phenotypic outcome vary dramatically as a function of the overall energy investment.

## Main

Living cells harness energy from the environment to drive out-of-equilibrium processes that promote and sustain life^7,8,11-14^. As nonequilibrium systems, cells extend classic thermodynamic principles, such as the second law (entropy production ΔS ≥ 0)^15,16^. Further, the complexity of subcellular processes poses a serious challenge in determining the rules that govern cellular metabolic energy expenditure *in vivo*^7,8^. For example, while recent studies measure cellular/organismal energy metabolism via calorimetry^17-20^, ATP fluorescence^21^, and metabolic or respiratory fluxes^9,22,23^, these techniques measure total energy, and cannot isolate the energy dissipated by concurrent internal processes that ultimately share the same source of energy. Consequently, how metabolic energy is used by and partitioned between non-equilibrium subsystems, and how this relates to biological outcomes remain enigmatic.

The eukaryotic cell cortex is a key driver of cell migration^24-28^, morphogenesis^4,5,29-32^, and cell division^3,33-35^, providing an ideal system to study energetic partitioning strategies at the intracellular level *in vivo*. The cell cortex comprises the plasma membrane and the underlying cytoskeleton, and itself can be considered to be composed of two subsystems. The first is a mechanical subsystem based on actin filaments (F-actin), the motor protein myosin-II, and other actin-binding proteins that perform mechanical work based on the consumption of ATP^36,37^. The second is a chemical signaling subsystem based on the G-protein Rho, which regulates F-actin and myosin-II by cycling between an active, GTP-bound form and an inactive GDP-bound form and thus directs cortical patterns based on the consumption of GTP. The interaction of these two systems makes the cell cortex a mechanochemical, dissipative, and excitable medium that can exhibit a variety of self-organized, periodic signaling, and mechanical patterns^2,3,23,32,38-40^, each associated with different phenotypic outcomes. For example, Rho and actomyosin pulses drive polarization in *C*.*elegans*^29^, and morphogenesis in *Drosophila* embryos^5^, and traveling waves of Rho and actomyosin drive cytokinesis in starfish and *Xenopus* embryos^3^. Here, we have exploited the periodic, quasi-two-dimensional nature of cortical pulses and traveling waves to study how energy is partitioned between the chemical and mechanical subsystems of the cell cortex under conditions where patterns are induced and systematically varied.

### Increased RGA-3/4 expression drives the cell cortex from quiescence through diverse excitable patterns

To systematically investigate how the cortex partitions energy between the chemical and mechanical subsystems, we expressed two cytokinetic regulators in immature *Xenopus* oocytes: Ect2, a protein that promotes the exchange of GDP for GTP by Rho and RGA-3/4 (aka ArhGAP11A or MP-GAP), a protein that promotes hydrolysis of GTP by Rho. The assembly of the F-actin is stimulated by Rho, while Rho is itself inactivated by F-actin. F-actin collaborates with myosin-II to transmit contractile force, triggering cortical deformation. Cortical deformation, in turn, impacts the chemical dynamics through advection. (The schematic is illustrated in Fig. 1A, B). The combined expression of Ect2 and RGA-3/4 converts the normally quiescent oocyte cortex into an excitable medium in which waves of Rho activity and “chasing waves” of F-actin and myosin-II develop^41^. While these waves mimic those observed in the cytokinetic furrow of dividing *Xenopus* and starfish embryos^3^, they are visible for many minutes (in contrast to furrow waves which rapidly ingress out of the reach of the microscope objective) rendering them amenable to quantitative analysis. Further, when the concentration of RGA-3/4 is varied against a background of constant Ect2 expression to modulate the consumption of GTP by Rho, a variety of periodic patterns are produced, ranging from pulses, to short wave-front choppy traveling waves, to labyrinthine traveling waves, to spiral waves (Fig. 1C, Movie S1, ref ^41^). The expected outcome of increased RGA-3/4 expression is not only an increase in the amount of energy consumed by the chemical subsystem of the cortex (via increased GTP hydrolysis by Rho) but also increased energy consumption by the mechanical subsystem. That is, while actin assembly and myosin-II consume ATP, actin assembly and myosin-II activation are gated by active Rho via formins and Rho-dependent kinase, respectively. Further, because RGA-3/4 expression increases cortical energy consumption, it is also expected to drive the cortex further from equilibrium, as will be established formally in the section below.

**Figure 1:**
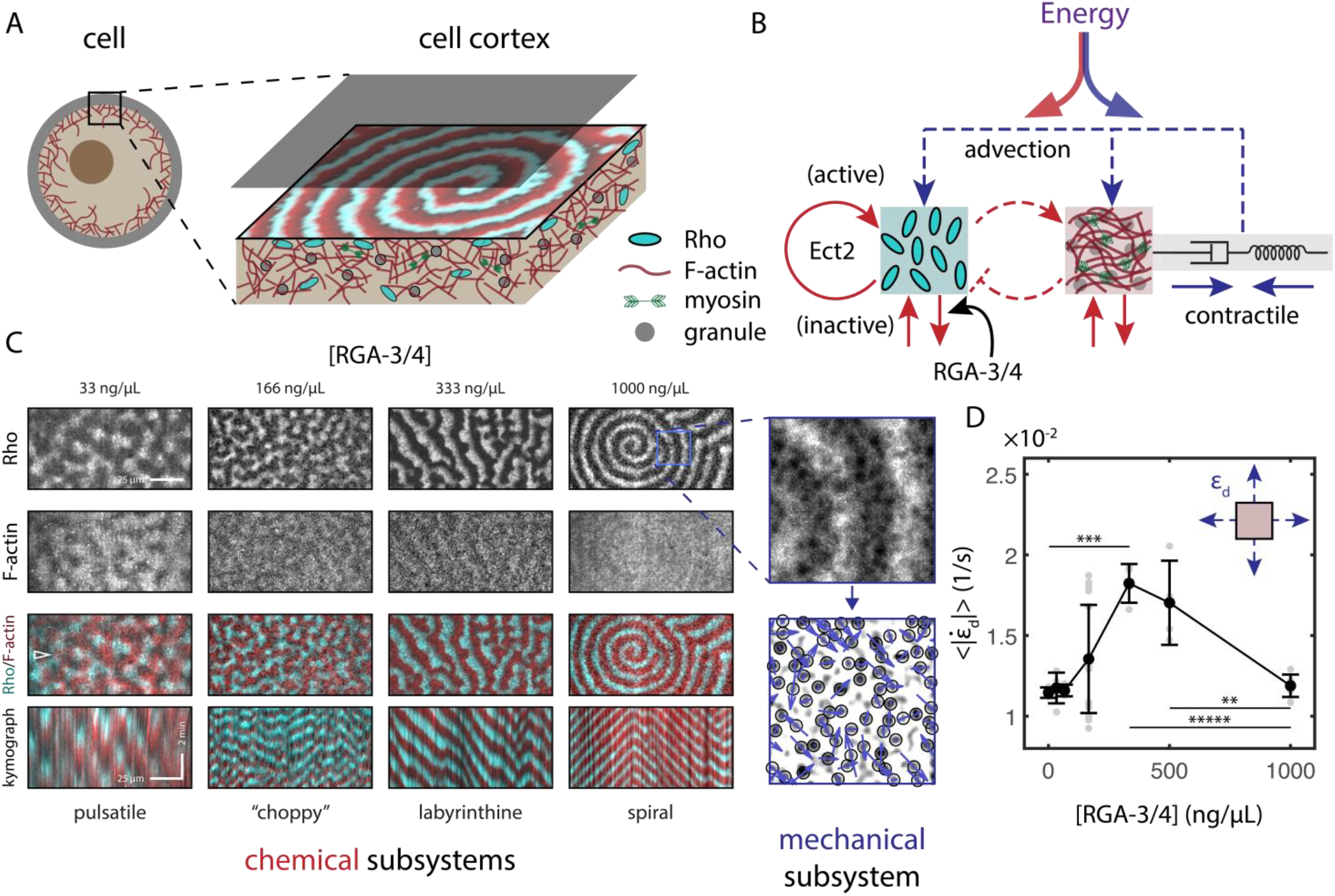
RGA-3/4 expression drives diverse mechano-chemical patterns in the cell cortex. **(A)** Schematic plot of the Rho/F-actin wave in the cell cortex. The spiral wave is an example from the simulation results with *k*_*RGA*_ = 0.6 1*/s*. **(B)** Schematic plot of the Rho/F-actin activator-inhibitor feedback loops and the coupling with the viscoelastic cell cortex. The red lines indicate the chemical reactions, and the blue lines indicate the coupled mechanical dynamics. The dashed lines indicate the interaction between two chemical subsystems (Rho and F-actin, red dashed lines), or between chemical and mechanical subsystems through advection (blue dashed lines). The arrow lines indicate promotion, and the T-shaped line indicates inhibition. Ect2 promotes the exchange of inactive Rho for active Rho, RGA-3/4 promotes the consumption of active Rho. **(C)** Experimental cortical system. Prototypical examples of pulsatile, choppy, labyrinthine, and spiral wave phenotypes with [RGA-3/4] = 33, 166, 333, 1000 ng/μL, respectively. From top to bottom, the rows show [Rho], [F-actin], a merged image with [Rho] and [F-actin] shown in cyan and red respectively, and a kymograph of the merged image (chemical signals). The kymograph is taken along a horizontal slice indicated by the open triangular symbol at the left of the merged image. Time is increasing downwards. Scale bars are 25 μm in space and 2 min in time. Inset on the right: zoomed-in portions from the blue box in panel (C) showing the pigment granules’ positions and their instantaneous velocities (blue arrows, the mechanical signal). **(D)** The absolute value of the dilation rate 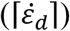 of the cortex deformation as a function of the [RGA-3/4].

The periodic cortical patterns were quantified by measurement of three signals, which relate to chemical and mechanical aspects of the periodic patterns: Rho (*R*, chemical) and F-actin (*F*, chemical/mechanical) were measured via fluorescence intensity (Methods). As shown by Fig. S4, both can be formulated as a noisy periodic oscillation with a deterministic part expressed by a periodic sinusoidal function, plus a noise part following the Gaussian distribution. The mechanical signal was quantified via local cell cortex deformation rate ( ***v***_*c*_, mechanical) and strain rates (dilation rate: 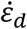, shear rate: 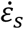, mechanical), based on the contraction-induced displacement of cortical pigment granules, which was previously shown to report myosin-powered bulk flow (Methods, Fig. S3, Movie S2)^42^. As previously described^41^, Rho wave amplitude initially increases sharply with increasing RGA-3/4 concentration ([RGA-3/4] from now on) but then reaches a plateau (Fig. S2). F-actin wave amplitude follows a similar pattern (Fig. S2). Curiously, however, after initially rising with increasing [RGA-3/4], the dilation rate, rather than plateauing, underwent a progressive decline back to the level obtained in the absence of RGA-3/4 (Fig. 1D).

To independently assess the consequences of experimental elevation of energy consumption by the chemical and mechanical subsystems, a computational model that combines signaling and mechanics via reaction-diffusion-advection equations with a coupled viscoelastic cortex material was developed (Methods). In this model, the increasing [RGA-3/4] is represented by increasing *k*_*RGA*_. This model faithfully captures the RGA-3/4-induced changes in two-dimensional cortical patterns seen in experiments (Fig. S1, Movie S3).

### RGA-3/4 expression determines energy partitioning in the cell cortex

We measured the entropy production rates (EPR, denoted as 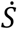) associated with progressively elevated [RGA-3/4]. EPR reflects the reversibility of system dynamics^43,44^ and is related to the dissipated energy. At steady state, the input energy is balanced by the energy output. The EPR represents a measure of the minimum energy required to sustain the observed process. Consequently, it offers a lower bound quantification to the input of energy, or the energetic cost, to sustain the periodic cortical patterns. The total EPR 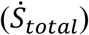 can be split into two nonnegative groups^45,46^: EPRs produced by each of the signals individually (Rho, F-actin, and granules’ deformation, giving 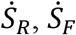 and 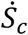) and EPRs produced by two interacting signals (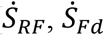 and 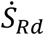). We take 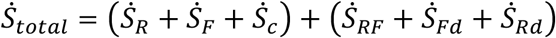. By investigating EPR generated by different chemical and mechanical dynamics, we are able to understand the energy partitioning among different subsystems (i.e.,*E*_*sub*_*/E*_*total*_) inside the cell cortex with the assumption that 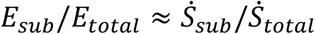.

First, we quantified 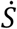 of each individual signal in the cell cortex by measuring the temporal autocorrelation for *R, F*, and ***v***_*c*_, independently. Autocorrelation functions of noisy oscillatory dynamics are typically given by sinusoidal functions oscillating with frequency *f* with an amplitude that decays exponentially with a time scale defined by the coherence time *τ* ^47-49^. 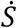 was estimated as 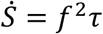 (Methods, wave coherence method). *f* (Fig. 2A) and *τ* (Fig. 2B) were extracted from the corresponding autocorrelation function (Methods, Fig. S5). We measured the 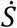 produced by chemical signaling from the Rho and F-actin dynamics (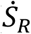 and 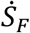), as well as by mechanical activities from the cortex deformation rate ***v***_*c*_, the dilation rate 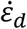, and the shear rate 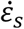 (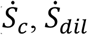 and 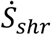). We find that the positive monotonicity of both *f* and *τ* at low [RGA-3/4] makes 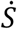 for all signals grow with [RGA-3/4]. At high [RGA-3/4], the positive monotonicity of *f* and the negative monotonicity of *τ* compensate, making 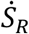 and 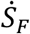 plateau, while in contrast, making 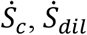 and 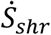 peak and decrease (Fig. 2C, Fig. S7). Overall, 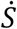 measured in simulation through autocorrelations and two other alternative methods align with the observed trends in experimental data. (Fig. 2F, Fig. S9, Methods).

**Figure 2:**
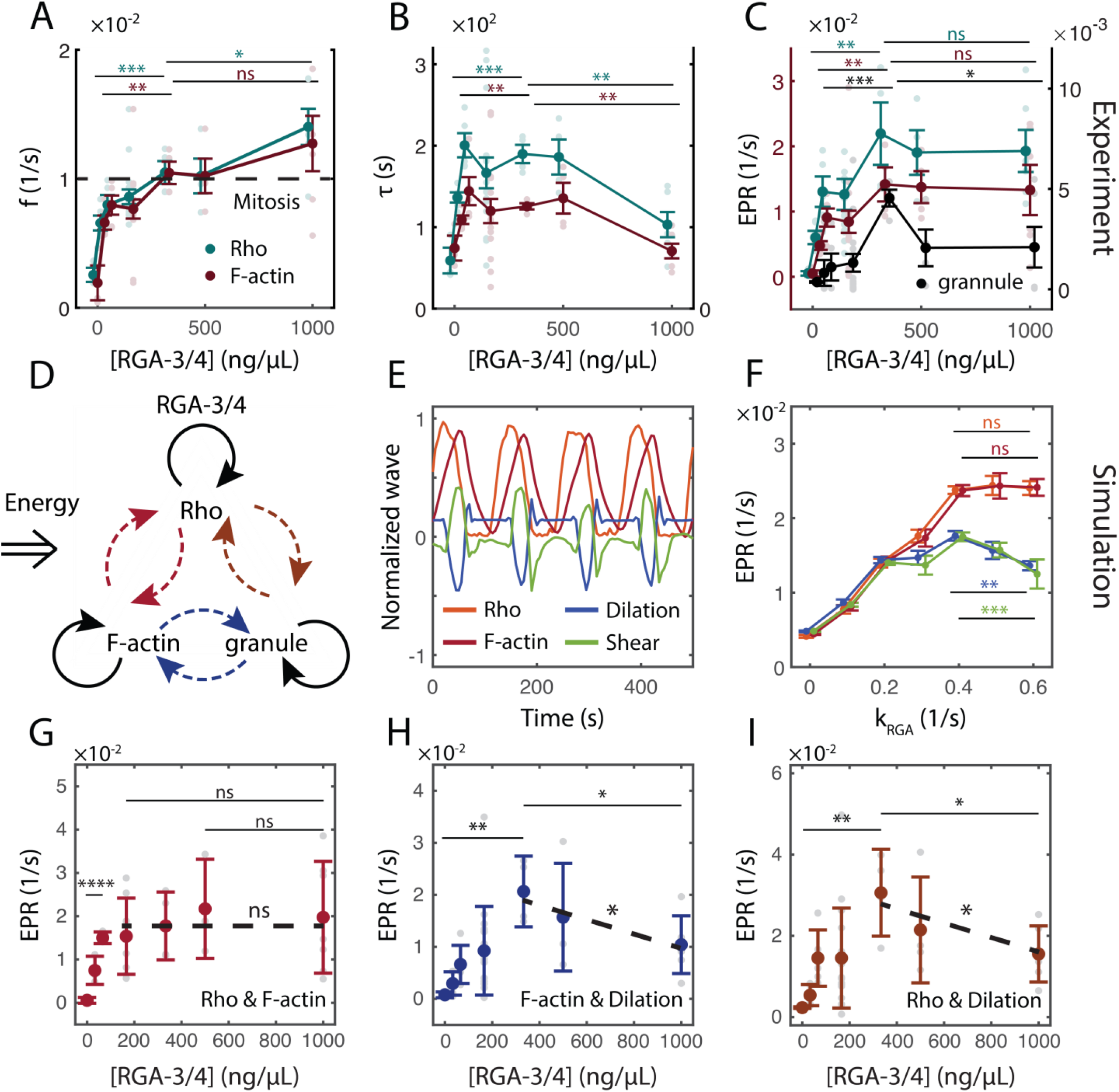
Energy partitioning between chemical and mechanical subsystems depends on RGA-3/4. **(A)** Autocorrelation frequency (*f*) as a function of RGA-3/4. **(B)** Autocorrelation coherence time (τ) as a function of RGA-3/4. **(C)** Entropy production rate, 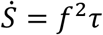, for Rho (cyan), F-actin (red), and granule motions (black). **(D)** Schematic plot of 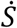 produced by autocorrelated and crosscorrelated signals. **(E-F)** Typical waveforms **(E)** of Rho (orange), F-actin (red), dilation rates (blue), and shear rates (green) in simulations and the corresponding 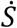 measured from the autocorrelations **(F). (G-I)** 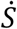 produced by interacting Rho & F-actin 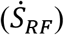 **(G)**, F-actin & dilation rates 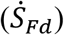 **(H)**, Rho & dilation rates 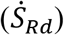 **(I)**. All statistical comparisons between two distributions were done with a two-sided *t*-test. We use the symbols *, **, ***, and **** for *p* < 0.1, 0.01, 0.001, and 0.0001, respectively. When fitting lines to data, we quote the *p*-value as significance values to rejections of the null hypothesis.

Comparing the quantifications (Fig. 2) to the patterns (Fig. 1) shows that as [RGA-3/4] increases, 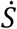 for the chemical subsystems grows as pulses give way to choppy traveling waves and then plateaus as choppy waves give way to labyrinthine and spiral traveling waves. In contrast, while 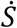 for the mechanical subsystem also initially increases in response to increasing [RGA-3/4], it drops at higher levels, corresponding to the point where choppy waves transition to labyrinthine traveling waves. Thus, interactions between the different components of the cell cortex qualitatively and quantitatively change as the energy consumed by Rho is progressively elevated.

### Decoupling of the chemical and mechanical subsystems at high RGA-3/4

The above results suggest that differential energy partitioning at high RGA-3/4 results in decoupling of the chemical and mechanical subsystems of the cell cortex. As an alternative way to test this inference, we used cross-correlation between each pair of *R, F*, and the cortex dilation rate 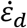 (Methods, Fig. 2D, Fig. S6). The 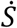 arising from *R* and *F* interactions, 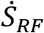 shows a similar trend as seen in the individual chemical signals, namely a monotonic growth that leads to a plateau (Fig. 2G). In contrast, the 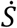 arising from both *R* and 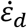 interactions, 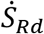 and *F* and 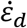 interactions, 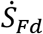, prominently peak and drop (Fig. 2H, I). These trends were replicated via a third method, namely, estimating the 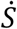 from interactions based on making a diffusive approximation for the dynamics^44^ (Methods, EPF method, Fig. S8), as well as in simulations (Fig. S9). Increased variance in the EPR is observed at high levels of [RGA-3/4] due to an increase in phenotypic variability and heterogeneity of the cortical waves (see Fig. S10D). Furthermore, the phase portrait analysis performed on the model corroborates the decoupling process (Supplemental Notes). As shown by Fig. 3A, with low *k*_*RGA*_, a limit cycle forms (which suggests periodic oscillations) only when the reaction-diffusion equation is coupled with the mechanical response. As a comparison in Fig. 3C, with high *k*_*RGA*_, a pure reaction-diffusion system can produce spontaneous Rho/F-actin oscillations that are little affected by the mechanical advective transport.

**Figure 3:**
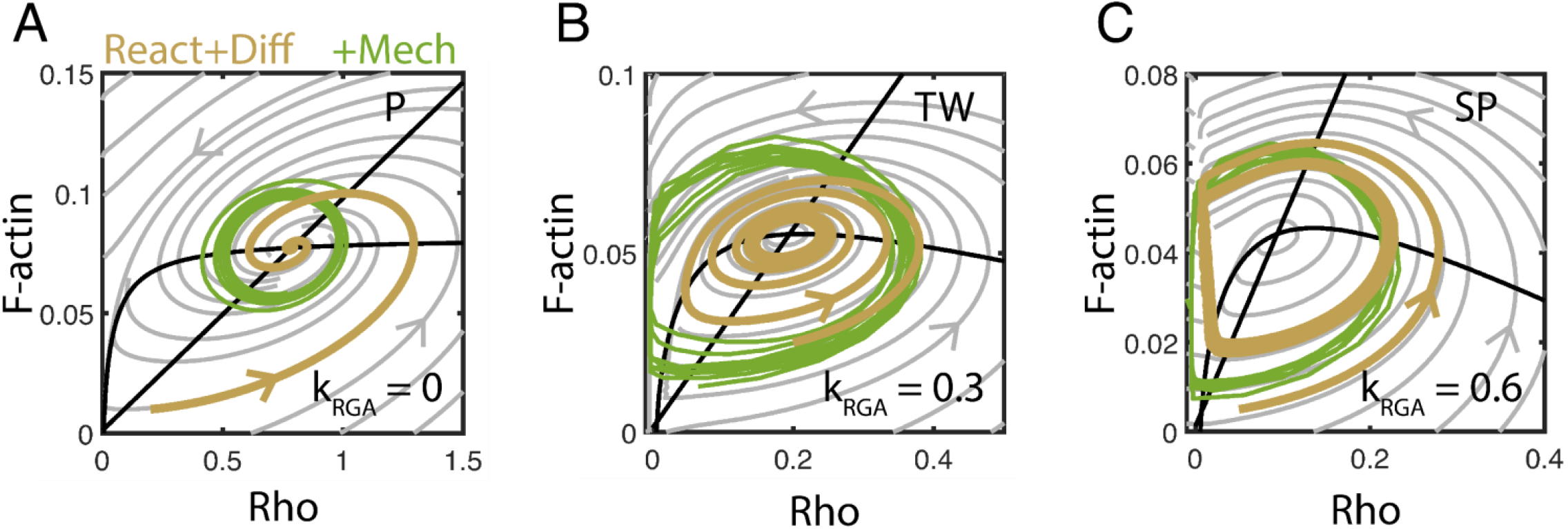
Mechanics get decoupled from chemical signaling in the highly activated cell cortex. Phase portrait analysis of different phenotypes in simulation: **(A)** Pulses (P). **(B)** Choppy traveling waves (TW). **(C)** Spirals (SP). Black lines: Nullclines of Rho and F-actin dynamical equations. Grey trajectories: a simplified system with only reaction terms (Model). Orange trajectories: a simplified system with only reaction-diffusion terms (Model). Green trajectories: a complete system with coupled reaction-diffusion-advection terms (Model).

Collectively, these results demonstrate that the energy input into the excitable cell cortex, controllable via [RGA-3/4], gets repartitioned to the chemical and mechanical subsystems of the cell cortex. At low [RGA-3/4], energy input leads to an increase in the 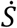 of Rho, F-actin, and mechanical deformations, while at high [RGA-3/4], the 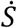 of Rho, F-actin dynamics and their interactions plateau, while the mechanical 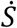 decreases, indicating the mechanical activities becomes decoupled from the chemical signaling.

### Onsager reciprocity suggests energy partitioning in a fixed proportion at modest RGA-3/4

Modestly elevating [RGA-3/4] makes mechanical 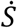 (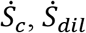 and 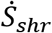) increase linearly with chemical 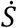 (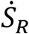 and 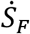), suggesting that, 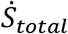 is partitioned among each subsystem in a fixed proportion. To test this inference and gain a deeper understanding of the inherent cross interactions amongst chemical and mechanical subsystems, we applied Onsager reciprocal relations, a theory that describes interactions between different pairs of fluxes (𝒥) and forces (ℱ) in subsystems^50^ (Fig. 4A).

**Figure 4:**
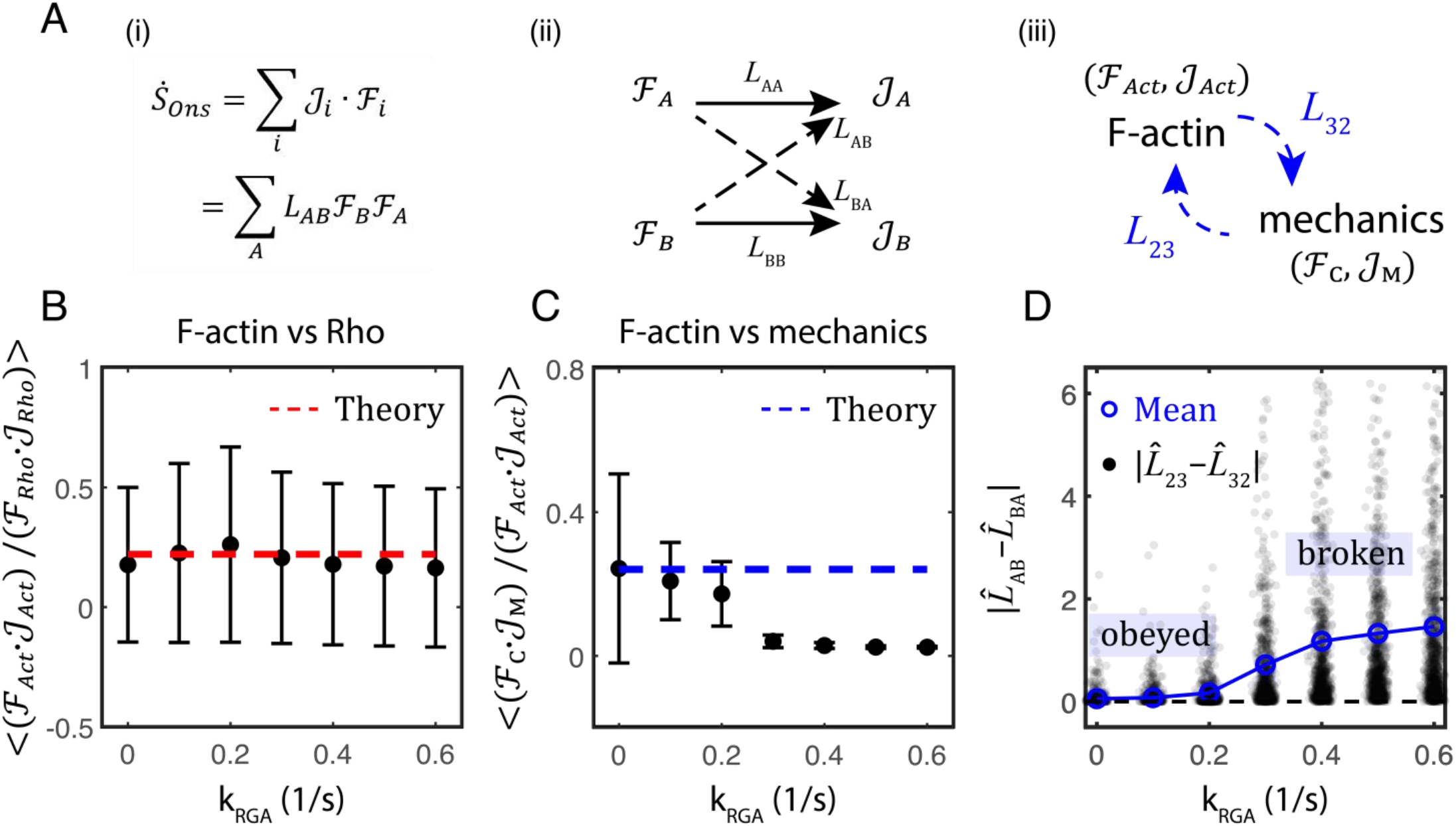
Onsager reciprocity implies energy partitioning among subsystems in a fixed proportion in simulation. **(A)** Equation and coefficients in Onsager’s theory: (i) expression of the total entropy production rate 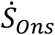, (ii) force-flux relationship, (iii) interactions between F-actin signaling and mechanical dynamics. **(B-C)** Ratio of the Onsager’s rate of entropy 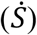 produced by F-actin dynamics (ℱ _*Act*_ ∙ 𝒥 _*Act*_) to that from Rho dynamics (ℱ *Rho* ∙ 𝒥 *Rho*) **(B)**, and mechanical dynamics (ℱ *C* ∙ 𝒥 *M*) to F-actin dynamics **(C). (D)** Absolute difference of the off-diagonal Onsager’s coefficients manifesting the F-actin/mechanics interactions. Data are normalized by *L*_22_ + *L*_33_. Open-colored circles are the mean values. Dashed lines are theoretical predictions. For each *k*_*RGA*_, n=1000.

We established the so-called Onsager matrix *L*^50,51^ (Methods). In Onsager’s theory, when the system is near equilibrium, there is a linear force-flux relationship^52-54^, based on which the Onsager matrix is constructed via 𝒥 = *L* ∙ ℱ with equal off-diagonal coefficients, i.e., *L*_*AB*_ = *L*_*BA*_ (*A* ≠ *B*), making *L* symmetric. In our case, we have three signals (*R, F*, and ***v***_*c*_), so *L* is a 3×3 matrix. To quantify Onsager reciprocity between each pair of ‘subsystems’, we measured the absolute difference |*L*_12_ − *L*_21_|, |*L*_23_ − *L*_32_|, and |*L*_13_ − *L*_31_|, which reflect the Rho/F-actin, F-actin/cortex motion, and the Rho/cortex motion interactions, respectively (Fig. 4A iii, Fig. S14). Measurements from both experiments (Fig. S10, Fig. S14) and simulations (Fig. 4D, Fig. S11, Fig. S14) demonstrate that with modest [RGA-3/4] (modest *k*_*RGA*_), |*L*_*AB*_ − *L*_*BA*_| ≈ 0, the pulsatile waves have a linear force-flux relationship (experiment: Fig. S12, simulation: Fig. S13) and the Onsager reciprocal relations are obeyed, as has been recently found for active systems in the linear response regime^55-57^. Strikingly, however, with large RGA-3/4 (or large *k*_*RGA*_), |*L*_*AB*_ − *L*_*BA*_| ≫ 0, the corresponding spiral waves have nonlinear force-flux relations (Fig. S12, Fig. S13) and broken Onsager reciprocal relations, as has been found with chiral active matter^58^. Interestingly, the choppy traveling wave occurs within but at the far end of the Onsager reciprocity (Fig. S10 D, Fig. S14) with linear force-flux relation (Fig. S12, Fig. S13).

To elucidate why energy partitioning among each subsystem in a fixed proportion at regimes with linear force-flux relation, we express 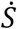 in Onsager’s form^50^ (Fig. 4A i). Here, 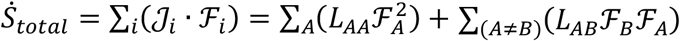, which decomposes the total EPR into contributions from individual subsystems (with label *A*), and interactions between pairs of subsystems (with label *A* and *B*). As shown clearly in the simulation (Fig. 4B-D) by the ratio of the Onsager’s rate of entropy 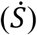 produced by F-actin dynamics (ℱ_*Act*_ ∙ 𝒥_*Act*_) to that from Rho dynamics (ℱ_*Rho*_ ∙ 𝒥_*Rho*_) (Fig. 4B), and mechanical dynamics (ℱ_*C*_ ∙ 𝒥_*M*_) to F-actin dynamics (Fig. 4C), the broken reciprocal relations initiate from where the differential energy partitioning occurs, suggesting that energy is partitioned among chemical and mechanical subsystems in a fixed proportion for wave dynamics with Onsager reciprocity. Theoretical derivation under the constraint of linear force-flux assumption verifies this observation (dashed lines in Fig. 4B, C, Supplementary Notes). Thus, we conclude that dynamic patterns in the cell cortex with Onsager reciprocity ([RGA-3/4] *≤* 166 ng/μL, *k*_*RGA*_ < 0.3 1*/s*) cause the partitioning of energy between the chemical and mechanical subsystems in a fixed proportion. Notably, we found an EPR “maximal” state at the far end of the Onsager reciprocity regime, represented by choppy wave dynamics wherein (i) the entropy production rate is maximized within the reciprocal regime, with (ii) strong coupling between chemical and mechanical subsystems as suggested by the coherence time measurement in Fig. 2B and phase portrait analysis in Fig. 3.

### Competing timescales determine the repartition of energy

Next, we investigated the energy partitioning between the chemical and mechanical subsystems beyond the range of Onsager reciprocity by taking advantage of the computational model. Here, we define two timescales: the chemical reaction rate *k*_*Chem*_ = *k*_*RGA*_, and the characteristic mechanical relaxation rate *k*_*Mech*_ = *A*_*s*_*/Gτ*_*ME*_ (Table I).

In the computational model (Model, Supplementary Materials), the chemical and mechanical subsystems are fully coupled by (i) actomyosin-induced contractile flow; (ii) mechanical properties of cell cortex affected by the F-actin concentration; (iii) the advective transport of chemical species. (ii) makes *k*_*Chem*_ a function of *k*_*Mech*_ (diamond symbols in Fig. 5A, B). To investigate the contribution of *k*_*Chem*_ and *k*_*Mech*_ to the energy partitioning independently, we decoupled *k*_*Chem*_ and *k*_*Mech*_ by making the shear modulus of the cortex (*G*) independent of the F-actin concentration. A phase diagram of decoupled *k*_*Chem*_ and *k*_*Mech*_ is shown in Fig. 5A, which captures all wave phenotypes observed in above sections. In Fig. 5B, we calculated the ratio of 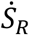 to the 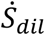, quantifying the proportion of the chemical dissipation over the mechanical dissipation. The results indicate that with fixed *k*_*Mech*_, increasing *k*_*Chem*_ leads to more energy partitioned to the chemical subsystems, and vice versa. Moreover, it predicts a regime where the mechanical dynamics gain more energy, in contrast to our findings in the above sections. This implies that the chemical and mechanical timescales compete in a manner that determines the partitioning of energy beyond the range of Onsager reciprocity.

**Figure 5:**
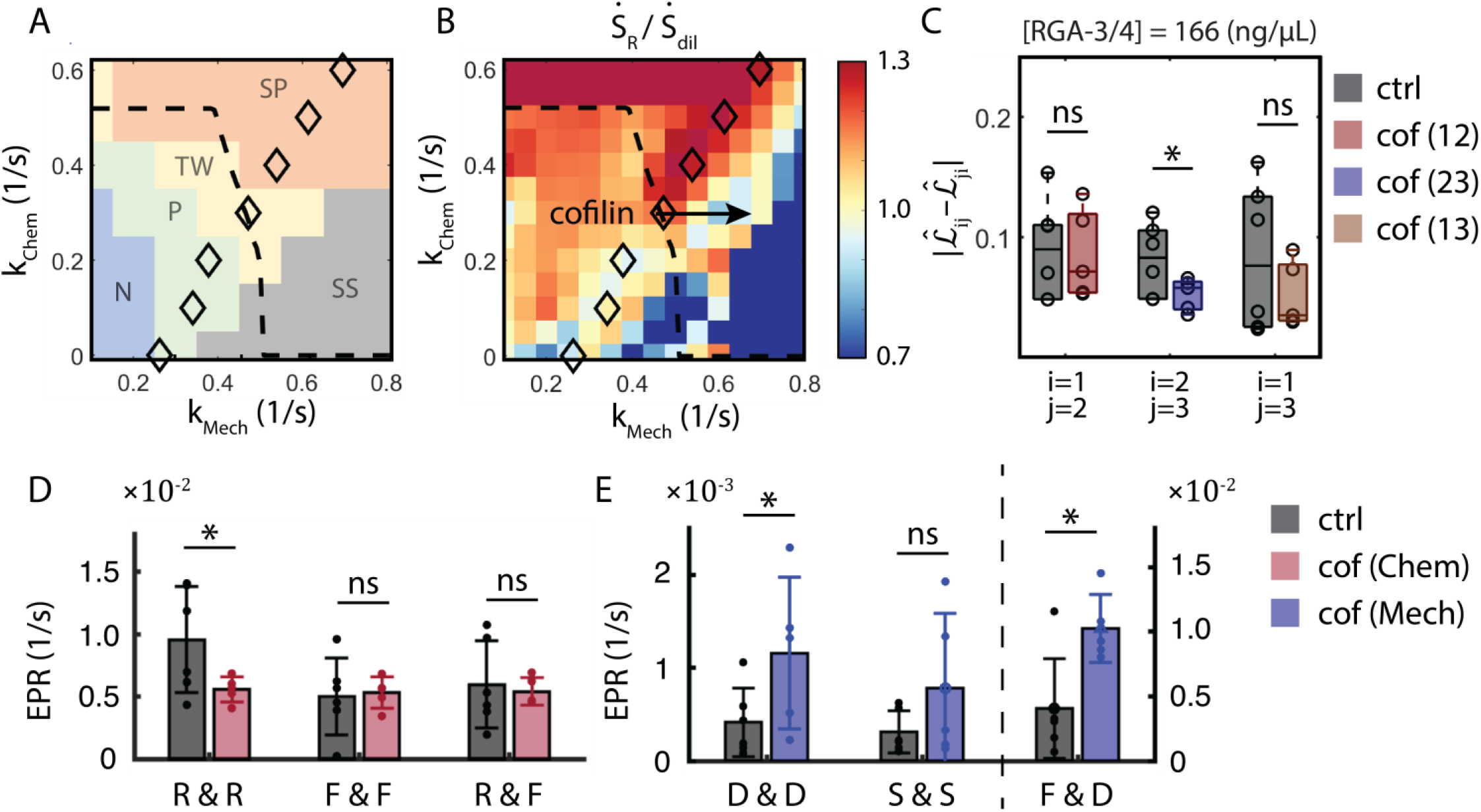
Competing internal timescales determine a repartition of energy. **(A)** Phase plane (*k*_*Mech*_, *k*_*Chem*_) with different color blocks classifying different wave phenotypes in simulation. N: noisy oscillations, P: pulses, TW: choppy or labyrinthine traveling waves, SS: self-reinforcing waves, SP: spirals. **(B)** Ratios of the EPR produced by Rho dynamics 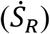 to the EPR produced by the dilation rates 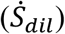 on the (*k*_*Mech*_, *k*_*Chem*_) phase plane. **(C)** Difference of the off-diagonal Onsager’s coefficients in cofilin experiments manifesting the Rho/F-actin interactions (red, *i* = 1, *j* = 2), F-actin/cortex motion interactions (blue, *i* = 2, *j* = 3), Rho/cortex motion interactions (orange, *i* = 1, *j* = 3). **(D)** EPRs are measured from the autocorrelations of Rho signals (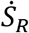, R&R), autocorrelations of F-actin signals (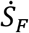, F&F), and cross-correlations of Rho/F-actin signals (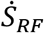, R&F). **(E)** EPRs are measured from the autocorrelations of dilation rates (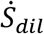, D&D), autocorrelations of shear rates (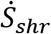, S&S), and cross-correlations of F-actin/dilation rates (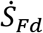, F&D). Grey bars mark the control experiments (n = 6), and colored bars mark the cofilin experiments (n = 5, red bars: chemical EPR; blue bars: mechanical EPR).

To experimentally verify this prediction, we performed experimental perturbations in which cofilin, an F-actin severing protein, was overexpressed *in vivo* to increase F-actin turnover (Methods). As a result, the mechanical subsystem can operate on a faster timescale to keep up with chemical subsystems, reminiscent of an increased *k*_*Mech*_ with roughly constant *k*_*Chem*_ in the model. Consequently, in cofilin experiments, restoration of the reciprocal relationship between F-actin dynamics and cortical motion that is caused by their more coordinated interactions is observed, when compared to control experiments (Fig. 5C). In Fig. 5D and 5E, we measured the 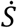 produced by the chemical and mechanical subsystems, respectively. This indicated that there is less chemical dissipation but more mechanical dissipation in the wave dynamics in cofilin experiments compared with the control experiments, verifying our theoretical predictions.

## Discussion

We manipulated energy consumption by the cell cortex via graded expression of the Rho GAP RGA-3/4 and measured the entropy production rate 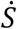 during steady-state wave behaviors. This measure reflects the minimum energy required to sustain the observed processes, not the actual energy, which also includes dissipation that takes place below optical resolution (due to ATP hydrolysis by actin and myosin and so on). By elevating GAP expression to drive the cell farther from equilibrium, we observed three sequential distinct regimes: (i) pulsatile wave with Onsager reciprocity, characterized by energy partition in a fixed proportion; (ii) choppy waves at the furthest extent of reciprocity also characterized by fixed energy partitioning; and (iii) labyrinthine or spiral traveling waves with broken Onsager reciprocity and characterized by differential energy partitioning between two subsystems. Notably, the wave frequency (*f*) monotonically increases with increased driving. In contrast, however, only within the Onsager reciprocity regime, displacing the system further from equilibrium increases 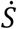 of both the chemical and mechanical subsystems in the cell cortex while maintaining strong couplings between them. This relationship between overall energy consumption and cortical dynamics culminates at the far end of this regime, where 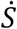 is maximized for both subsystems and is characterized by choppy waves with intermediate *f*. Intriguingly, this state of maximal 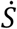 bears a striking resemblance, both qualitatively and quantitatively (with a frequency of approximately 0.01*s*^−1 10^), to the cortical waves associated with cell division^3,10,41^. We speculate that this EPR maximal state within the reciprocity regime is particularly advantageous for cell division, as it ensures both a high level of chemical/mechanical activities and chemical-mechanical coupling, potentially facilitating the robust functioning of cell division. A similar observation was also identified in crawling Dictyostelium cells^59^, where actin oscillations associated with cell motility and chemotaxis were induced by a signaling stimulus. The most excitable state occurs neither with the most rapid nor the slowest stimulus, but at an intermediate state near the instability border where the signaling pathway and actin responses are well coordinated. In stark contrast, a departure from this state at high levels of GAP expression with broken Onsager reciprocity results in a differential energy partitioning among the now-uncoupled subsystems, where the mechanical activities struggle to keep pace with the excessively rapid chemical oscillations. It is essential to emphasize that such an uncoupled state can pose severe threats to the cell’s viability and function^60^. For instance, impaired excitation-contraction coupling in cardiac muscle, leading to a loss of synchronization between electrical and mechanical activities, is known to be a potentially life-threatening condition that culminates in sudden cardiac death^61^.

Our results reveal fundamental principles in cellular energetics and nonequilibrium physics governing the intricate interplay of energy partitioning and subsystem coupling in cellular dynamics. Furthermore, the underlying physics of cortical excitability clearly extends beyond embryonic cells^40^, suggesting that our results and their implications will be relevant across diverse cell types.

## Methods

Methods and any associated references are available in Supplementary Materials.

## Contributions

M.P.M. and W.M.B. designed and conceived the project. W.M.B. designed and conceived the experimental work. M.P.M., S.C., and D.S.S. designed and conceived the analytical, computational and theoretical work. W.M.B., A.M., and S.K. performed experiments. S.C. and D.S.S. analyzed experimental data. S.C. implemented the model, performed and analyzed simulations. S.C. did the theoretical analysis. D.S.S. did the theoretical analysis of the Stuart-Landau equation. S.C., D.S.S., W.M.B., and M.P.M. wrote the manuscript.

## Notes

### Competing Interest Statement

The authors have declared no competing interest.

